# Transient CRISPR immunity leads to coexistence with phages

**DOI:** 10.1101/2019.12.19.882027

**Authors:** Sean Meaden, Loris Capria, Ellinor Alseth, Ambarish Biswas, Luca Lenzi, Angus Buckling, Stineke van Houte, Edze R Westra

## Abstract

Phages play a major role in shaping the composition, evolution and function of bacterial communities. While bacteria and phages coexist in many natural environments, their coexistence is often short-lived in the lab due to the evolution of phage resistance. However, fitness costs associated with resistance and mutational loss of resistance alleles may limit the durability of acquired resistances, potentially allowing phages to re-invade the population. Here, we explore this idea in the context of bacteria that evolve CRISPR-based immunity against their phages. Consistent with previous studies, we found that the bacterium *Pseudomonas aeruginosa* PA14 evolved high levels of CRISPR-based immunity and low levels of surface-based resistance following infection with phage DMS3vir, which led to rapid phage extinction. However, when these pre-immunized bacterial populations were subsequently challenged with the same phage, they failed to clear the infection and instead stably coexisted with the phage. Analysis of bacterial genotypes and phenotypes over time explained why CRISPR-Cas immunity provides only a transient advantage: in the absence of phage (i.e. following the initial phage extinction) formerly CRISPR-immune bacteria regain sensitivity due to evolutionary loss of spacers, whereas in the presence of phage (i.e. upon reinfection) selection favours surface-based resistance over CRISPR immunity. The latter results from an infection-induced fitness cost of CRISPR-immunity that is due to phage gene expression prior to target DNA cleavage by the immune system. Together, these results show that CRISPR-Cas immune systems provide only a transient benefit to bacteria upon phage infection and help to explain why bacteria and phages can coexist in natural environments even when bacteria carry CRISPR-Cas adaptive immune systems that allow for rapid acquisition of immunity against phages.

## Main

Phages are ubiquitous in prokaryotic communities and play a vital role in key ecological processes. For example, phage predation may cause up to 20% of mortality of all bacteria in the world’s oceans, making them key players in global nutrient cycling (Suttle 2007). In the deep terrestrial biosphere, phages regulate microbial communities- in turn shaping resource turnover (Daly *et al.* 2019) and in wetland ecosystems, phages have been associated with regulation of methanogens and sulfate reducers (Dalcin Martins *et al.* 2018). In addition to ecological processes, co-evolution between phages and bacteria can generate and accelerate increases in bacterial and phage genetic diversity (van Houte *et al.*; Paterson *et al.* 2010; Scanlan *et al.* 2015), mediate horizontal gene transfer (Canchaya *et al.* 2003) and alter competition among bacteria (reviewed in Koskella & Brockhurst 2014). As well as this important ecological role, phages are also increasingly seen as a promising alternative to antibiotics to treat bacterial infections of plants, animals and humans (Wagenaar *et al.* 2005; Balogh *et al.* 2010; Kutter *et al.* 2010). Therefore, understanding the factors that shape the ecology and evolution of bacteria-phage interactions is fundamentally important.

The impact of phages on bacterial species and communities in ecological and biomedical settings is inevitably constrained by both pre-existing (innate) and acquired phage resistance (reviewed in Labrie *et al.* 2010). Bacteria can acquire phage resistance either by mutation or masking of the cell surface receptors to which the phages bind, or through their adaptive immune systems known as CRISPR-Cas (Clustered Regularly Interspaced Short Palindromic Repeats; CRISPR-associated). CRISPR-Cas works by inserting phage-derived sequences into CRISPR loci on the host genome, which function as genetic memory to detect and destroy phages carrying the same sequence (reviewed in Marraffini 2015). This can lead to rapid phage extinction, as phages are effectively being actively removed from the environment (as opposed to surface-based resistance where phages persist in the environment).

While CRISPR immunity can drive phages extinct in the short term, it is less clear how this immune system impacts phage persistence in the long term. Given that acquired resistance alleles are heritable, one would perhaps expect that pre-immunized bacterial populations would remain fully resistant to reinfection by the same phage genotype. On the other hand, there may be selection against CRISPR immunity due to costs in the form of autoimmunity (Stern *et al.* 2010), or if CRISPR is less effective under high phage exposure (Westra *et al.* 2015). Additionally recent theory and data suggest that individual bacteria may lose their CRISPR-resistance due to mutations in the *cas* genes or CRISPR arrays (Weissman *et al.* 2019).

To test the durability of CRISPR-based immunity, we recurrently infected *Pseudomonas aeruginosa* PA14 with phage DMS3vir (Cady *et al.* 2012), a mu-like phage (Budzik *et al.* 2004). By sequentially re-introducing the phage at different time intervals we vary the intensity of selection for resistance (surface mutation) and immunity (CRISPR) and track the resulting ecological dynamics. By using a combination of experimental evolution and genomic analyses we tease apart the population genetics that underpin this host-parasite interaction. These results have implications for how genetic diversity can be both generated and lost in bacterial populations and help explain the coexistence of bacteria and phages in natural microbial communities.

## Results

### Populations lose the ability to drive phages extinct

CRISPR mediated immunity acts as an acquired, heritable genetic memory against phages (Marraffini 2015), however the timescales over which it is effective are unknown. Based on our mechanistic understanding of CRISPR immunity, one would predict that population-level immunity is long-lived. However, the few empirical studies that directly measure population and evolutionary dynamics of CRISPR-phage interactions have either focussed on short-term dynamics (Westra *et al.* 2015; van Houte *et al.* 2016) or examined CRISPR in isolation (Paez-Espino *et al.* 2013; Common *et al.* 2019), rather than the interplay between alternative resistance strategies. Recent theory and data suggests that resistance alleles may be lost from bacterial populations due to back mutations towards a sensitive phenotype when resistance is costly, which can in turn allow for phage persistence (Weissman *et al.* 2018; Chaudhry *et al.* 2018; Gurney *et al.* 2019). To test this, we exposed *P. aeruginosa* PA14, which carries a type I-F CRISPR system, to phage DMS3vir, which carries a mutation in its repressor gene that prevents lysogeny, as well as a partial protospacer match which promotes primed spacer acquisition by *P. aeruginosa* PA14 (Westra *et al.* 2015). We then measured whether populations could repeatedly drive phage DMS3vir to extinction following an initial phage exposure. Consistent with earlier findings, following the first infection treatment, phage titers were found to initially increase due to the phage epidemic that takes place in the sensitive bacterial population, followed by a rapid decline in phage titers and extinction at 4 days post infection (dpi) (Fig. 1A), due to the evolution of CRISPR-based immunity in the bacterial population (van Houte *et al.* 2016 and Fig. 2). To test whether these now immunized bacteria could survive subsequent infections they were reinfected with 10^7^ pfu phage at 5 dpi. Consistent with the bacterial population having evolved CRISPR-mediated phage resistance in response to the earlier phage challenge the phage was rapidly driven to extinction (Fig. 1B). Strikingly, when the same bacterial populations were re-infected 5 days later (at 10 dpi) they had lost the ability to drive the phage extinct (Fig. 1B), and phage was able to persist for the remainder of the 30-day experiment in all replicates of the same treatment. Similarly, in the treatment where the first re-infection took place at 10 dpi, the bacterial populations also had lost the ability to drive phage extinct (Fig. 1C). As expected, the treatment where phage was added every day showed high phage titres throughout the experiment (Fig. 1D), and in the phage-free treatment no phage could be detected (data not shown).

**Figure 1.**
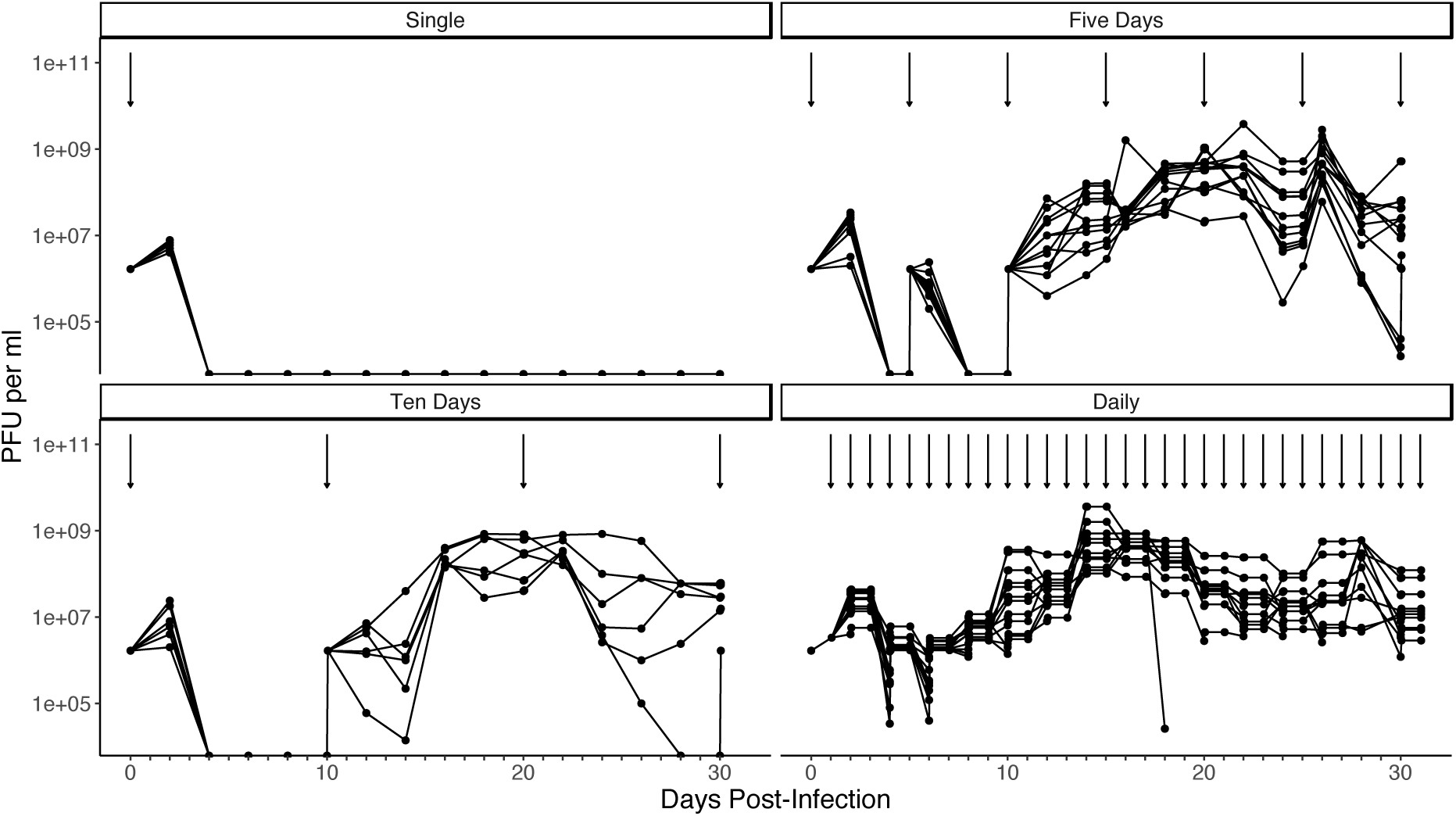
Phage titres over time upon infection of *P. aeruginosa* with DMS3vir. Arrows denote timing of reinfection, each line represents an individual replicate population. A) No reinfection (n = 6). B) Reinfection every 5 days (n = 12). C) Reinfection every 10 days (n = 6). D) Daily reinfection. (n = 12). Phage titres were recorded every 2 days. Note this figure includes imputed values for non-recorded days that combine the previous days titre with the phages that were added. Limit of detection is 200 PFU per ml.

**Figure 2.**
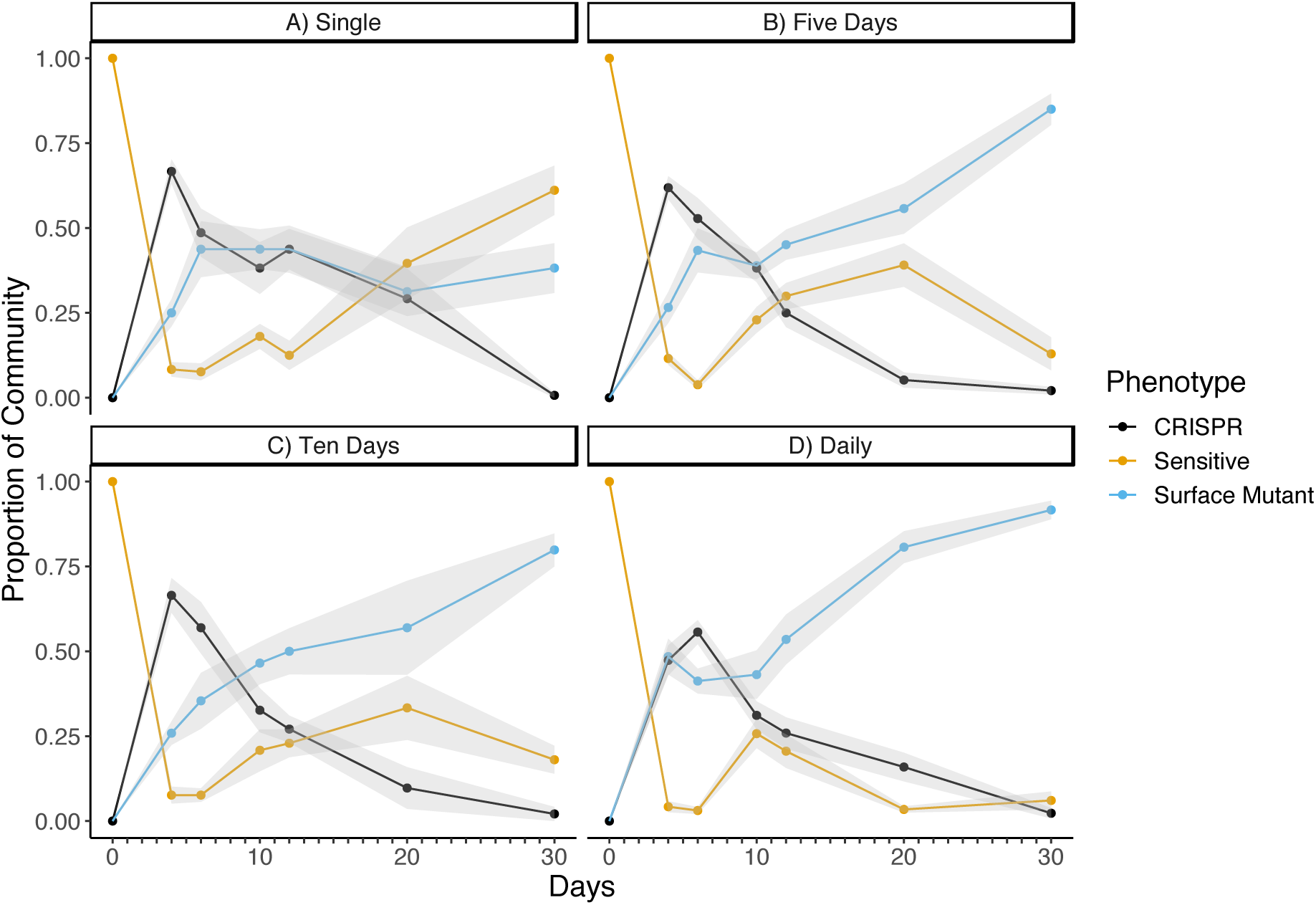
Frequencies of phage resistant and sensitive phenotypes. A) No reinfection, n = 6 B) Reinfection every 5 days, n = 12 C) Reinfection every 10 days, n = 6, 5D) Daily reinfection, n = 12. Proportions were determined from phenotypic assays of 24 clones per population. Lines represent the means across populations and shaded areas represent ±1 SEM. CRISPR-immune bacteria are shown in black, bacteria with surface-based resistance in blue, and clones that remain sensitive to the phage in yellow.

### Populations are invaded by surface resistance mutants and sensitive clones

To understand why phages can successfully re-infect populations that previously drove the same phage extinct, we phenotypically characterised the composition of the host populations, i.e. the proportion of sensitive bacteria as well as the proportion of bacteria that had acquired at least one new spacer that conferred CRISPR-based immunity (CRISPR-immune herein) or surface mutation based resistance (i.e. bacteria that evolved phage resistance through loss or masking of the phage receptor, SM herein). At 4 dpi 0.66 ± 0.036 (mean proportion ± SE) of bacteria in the populations from the single infected treatment evolved CRISPR-immunity, 0.25 ± 0.040 evolved SM resistance, and 0.083 ± 0.022 remained sensitive. At 30 dpi, these fractions had changed to 0.0069 ± 0.0069 CRISPR immunity, 0.38 ± 0.073 SM, and 0.61 ± 0.73 sensitives. The invasion of sensitive bacteria in the absence of phage suggests that either CRISPR immunity is associated with a constitutive fitness cost, or that there is back mutation from CRISPR immunity to the sensitive, ancestral genotype (examined in detail later).

Next, to understand how the reinfection regimes impact the evolutionary dynamics of the host population, we examined how the evolution of CRISPR immunity and surface resistance varied across our treatments. At 4 dpi, we found an interaction between the proportion of each phenotype and reinfection frequency (GLM, F1, 102 = 7.06, p < 0.0001), with populations that were re-infected daily had slightly increased levels of SM compared to those only infected once (F_1,35_ = 17.94, p < 0.001), which is consistent with previous results that demonstrate a shift towards SM when phage exposure is high (Westra *et al.* 2015). Interestingly, for all treatments, we observed a significant reduction in CRISPR-immune clones over time (quasibinomial GLM, F_1,22_ = 179.8, p < 0.0001), but no interaction between treatment and time (quasibinomial GLM, F_1,16_ = 2.4, p = 0.16), suggesting the reinfection regime was not driving the observed decline. However, while in the single infection treatment the decline in CRISPR-immune clones was due to the invasion of sensitive bacteria, we observed invasion of surface resistant bacteria in the reinfection treatments instead (Fig. 2).

### The population genetics of phage resistance and immunity

These phenotypic data were further supported by genotypic data. First, we sequenced the genomes of a single SM clone from each population at day 12. Consistent with phage DMS3vir being pilus-specific (Budzik *et al.* 2004), we identified in all cases a SNP in a pilus affiliated gene (PilC, PilB, PilQ, PilY1), a deletion of the PilM gene or, most frequently, a 10kb deletion of 7 pilus-associated genes (at position 5948050 – 5958338; Fig. 3, panel E), which was identical in all independent isolates and hence suggests these areas of the genome contain sequences with microhomology that are prone to recombination. In addition, we carried out deep sequencing of CRISPR array amplicons (CRISPR1 and CRISPR2) of all experimental populations. This demonstrated an extremely high spacer diversity (133249 unique CRISPR arrays) that is sufficient to cover all 5643 possible target sequences on the phage genome (Fig.S1). We found a significant interaction between time and reinfection frequency, with spacer diversity declining in all groups with the exception of populations that were infected daily, which maintained spacer diversity for longer (GLM, F_1,85_ = 6.65, p < 0.001, with qualitatively similar results for Shannon’s or Simpson’s diversity indices). Moreover, the frequency of array sequences lacking any new spacers increased over time (GLM, F_1,89_ = 228, p < 0.0001), as would be expected given the invasion of sensitive and SM clones. Interestingly, arrays with multiple spacers all declined throughout the experiment (up to 5 spacers, Fig 3, Panel A). When we tracked the fates of individual spacers (in order to identify variation in spacer effectiveness) we typically found a consistent decline in spacer abundance, with only a few exceptions (Fig 3, panel B) that most likely represent CRISPR immune clones that subsequently acquired a surface mutation (Fig. 3, Panel C).

**Figure 3.**
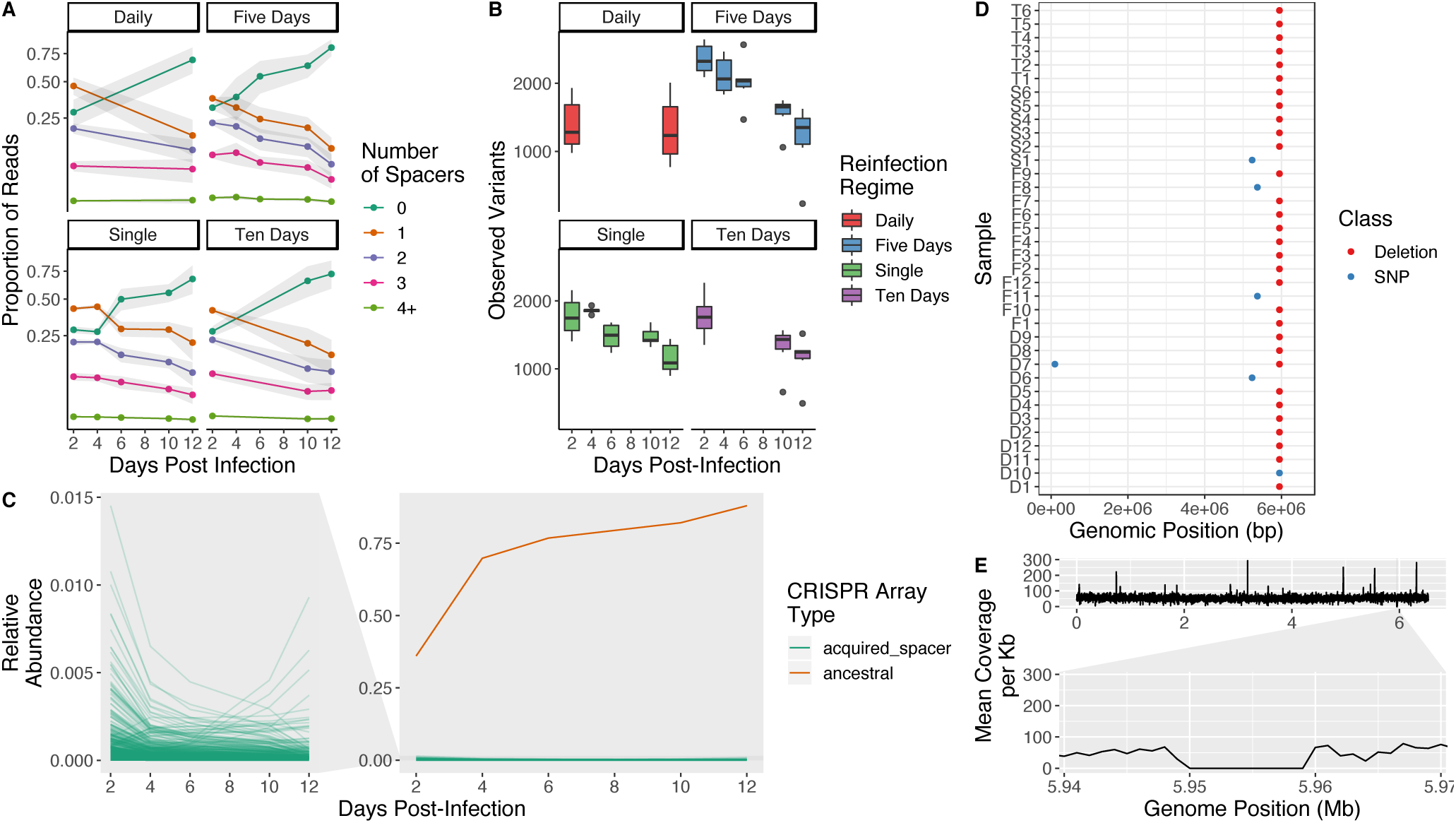
Population genetics of CRISPR arrays and surface mutants. A) Mean proportion of reads with acquired spacers per treatment. Ribbon denotes 1 S.D. B) Decline in genetic diversity in spacer arrays across treatments. These results are qualitatively similar regardless of diversity index used (i.e. Shannon, Simpson’s). C) Frequencies of individual arrays through time for a single population of the five-day reinfection regime. Orange line represents the ancestral array, with no new spacer. Green lines represent unique genotypes. Zoomed panel shows individual spacer frequencies. Figures show the results for CR2, the more active locus, however these results are also representative of CR1. D) Whole genome analysis of re-sequenced clones exhibiting a surface modification phenotype at day 12-points denote SNP locations. Sample names denote infection regime and replicate. E) Mean coverage of reads across the PA14 genome, for a single sample (S2), split by 1Kb intervals. Zoomed panel shows the 10,288 bp deletion containing pilus genes.

### Sensitive bacteria act as a reservoir for phage amplification

The data discussed above show that CRISPR immunity dominates early on, but over time sensitive bacteria and SM clones invade. Since sensitive bacteria will act as a source for phage, we hypothesised that their invasion may be responsible for the observed bacteria-phage coexistence following re-infections at 10 dpi. Consistent with this hypothesis, bacterial populations from 2 dpi were able to rapidly reduce phage DMS3vir titres whereas the same populations from 12 dpi caused phage amplification (GLM, F_1,46_ = 172.29, p < 0.0001, Fig. S2). Moreover, we found a strong interaction effect between phage titres and the proportion of each phenotype in our experimental populations (GLM, F_1,627_ = 50.93, p < 0.0001). Specifically, we found a positive correlation between phage titre and the frequency of sensitive clones in the population (GLM, F_1,209_ = 12.3, adjusted p-value < 0.001), as well as a negative correlation between phage titre and the frequency of CRISPR-immune clones (GLM, F_1,209_, F = 41.6, adjusted p-value < 0.001) (Fig 4.). As expected, the proportion of clones with SM resistance correlates less strongly with phage titre (as this phenotype neither amplifies nor degrades phages). Taken together, these results are consistent with the idea that CRISPR clones act as a ‘sink’ for phages whilst sensitive bacteria act as a ‘source’ for phages and that the protective effect is lost as CRISPR-immune bacteria decline in frequency.

**Figure 4.**
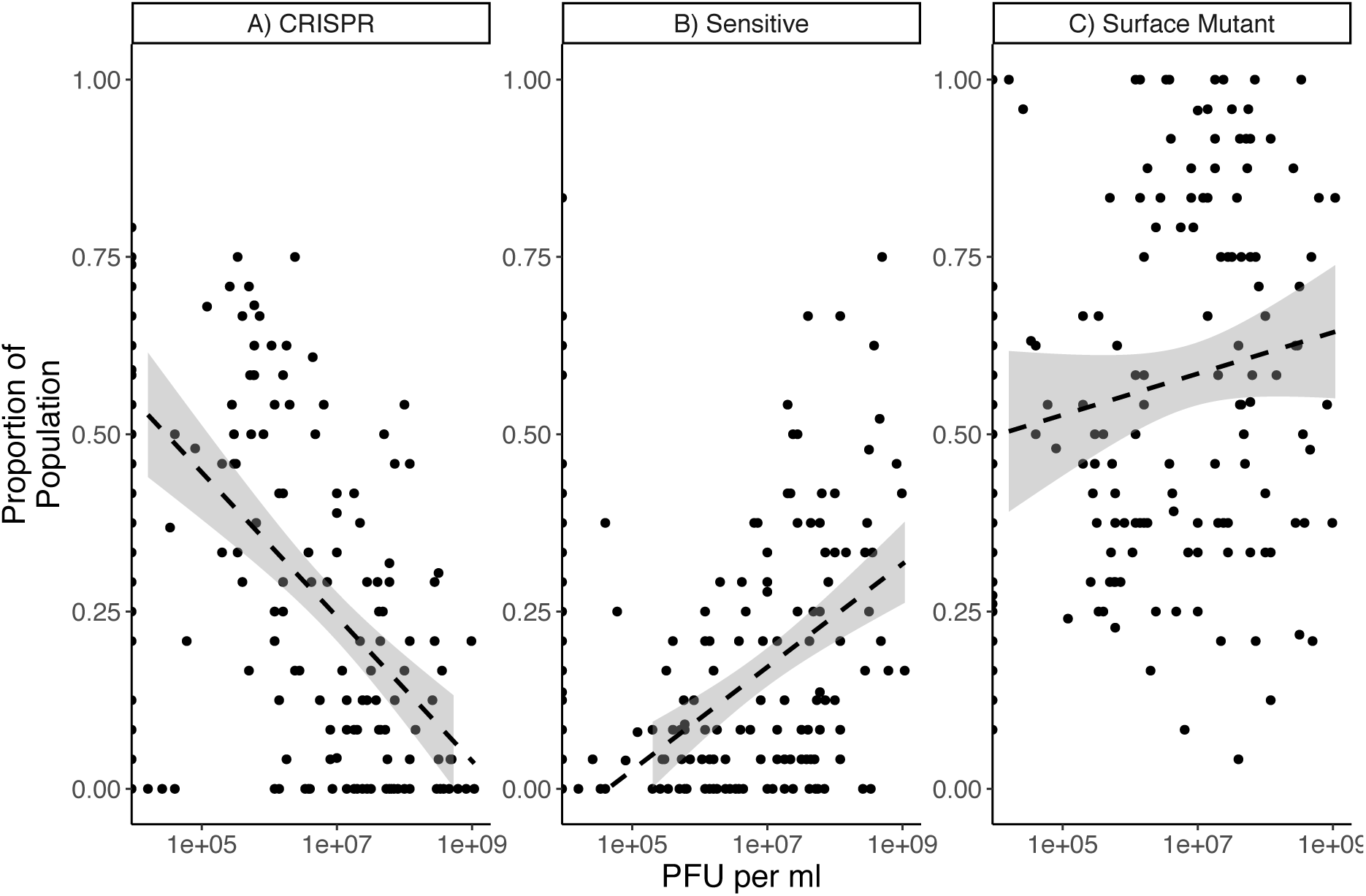
Correlations of each phenotype with phage titres throughout the experiment. Dashed lines represent a linear model fit and shaded areas represent 95% confidence intervals. A) Surface mutant bacteria, B) CRISPR-immune bacteria, C) Sensitive bacteria.

### Phage densities drive the host evolutionary dynamics

Whereas invasion of sensitive bacteria can explain the phage population dynamics, it is less clear what drives the invasion of sensitives in the first place. To understand this, we first compared the host dynamics across the different infection regimes. Even though the decline of CRISPR-immune clones was broadly similar across the various infection regimes, there were clear differences in the invasion of sensitive bacteria and clones with SM resistance that depended on the phage titers: sensitive bacteria were most likely to increase in frequency when phages were absent, whereas SM tended to invade in the presence of phage (GLM, interaction between phage presence and phenotype on change in frequency, F_1,611_ = 30.87, p < 0.001, Fig S3). Invasion of sensitives in the absence of phage may be driven by selection, if sensitive bacteria are fitter than clones with CRISPR immunity under those conditions. However, the observed increase in the frequency of sensitives may also result from back-mutation from a CRISPR immune genotype to sensitive (for example mutations in *cas* genes can inactivate the immune system (Jiang *et al.* 2013) or recombination between CRISPR repeats can result in spacer loss (Lopez-Sanchez *et al.* 2012). To tease these processes apart, we competed clones with different phenotypes isolated from these evolution experiments against an ancestral reference strain in the absence of phages. We found that although the clones did increase in fitness by day 12 (GLMM, main effect of time, χ^2^ = 31.07, df = 6, p < 0.001, Fig. 5), there was no interaction between phenotype and time (GLMM, χ^2^ = 3.11, df = 11, p = 0.21) and no effect of treatment (GLMM, χ^2^ = 1.83, df = 9, p = 0.61). We also found a borderline significant effect of phenotype (GLMM, χ^2^ = 6.54, df = 6, p = 0.038), however post-hoc testing showed that there were no significant differences among phenotypes within each timepoint (Table S1). When considered alongside the observed ecological dynamics, the most parsimonious explanation is that sensitive bacteria invade the populations through the loss of CRISPR immunity i.e. back-mutation where spacers are lost. This is most clear in the single infection treatment where phages are absent: no phenotype has greater fitness yet sensitive bacteria still invade (Fig. 2).

**Fig. 5.**
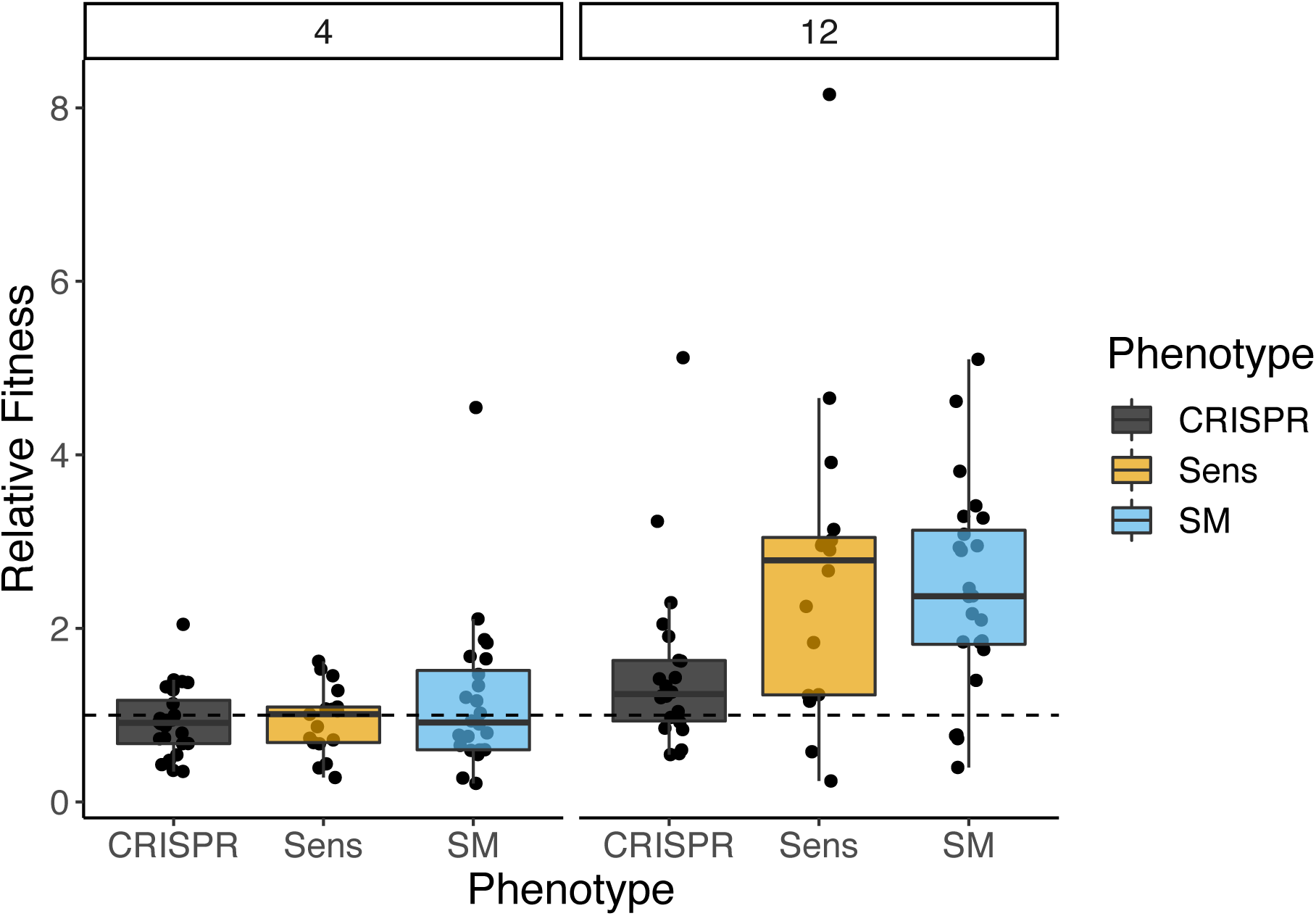
Relative fitness of bacterial clones at early (day 4) and late (day 12) timepoints during the experiment (in the absence of phages). Relative fitness was determined from 3-day competition experiments against a marked reference strain in the absence of phages. Where possible, a clone of each phenotype was isolated from all populations. A) Clones isolated from 4 DPI, B) Clones isolated from 12 DPI.

In contrast, the rapid invasion of SM clones observed in the evolution experiment, in the presence of phage, is likely explained by the fact that CRISPR immunity is associated with an inducible cost (i.e. a cost that increase during phage infections (Westra *et al.* 2015). Indeed, a similar invasion of SM has been previously observed when a reservoir of sensitive bacteria was introduced because the force of infection is increased (Chabas *et al.* 2016). However, it has remained unclear what the mechanistic basis of this cost is. We envisaged three possible reasons for this inducible cost: (i) a cost of autoimmunity, where bacteria acquire self-targeting spacers during infections (ii) a fitness cost due to enhanced expression of the CRISPR-Cas immune system following infection or (iii) due to expression of phage-encoded genes in the period between infection (i.e. phage genome injection) and CRISPR-mediated clearance of the phage. To explore the first hypothesis, we first performed a competition experiment between bacteria with CRISPR immunity that are unable to acquire novel spacers (i.e. a *cas1* deletion mutant) and a surface mutant. This competition was performed across a gradient of titres of DMS3mvir (a mutant of DMS3vir that is targeted by spacer1 in CRISPR array 2 of the PA14Δ*cas1* strain; hence both the surface mutant and competing bacterial genotypes being insensitive to this phage). If the induced fitness cost of CRISPR-Cas immunity was due to the acquisition of self-targeting spacers, we would expect the *cas1* deletion mutant to outcompete the WT strain. However, this was not observed, with the *cas1* deletion mutant showing lower relative fitness than the WT strain (GLM, F_1,70_ = 17.85, p < 0.001, Fig. S4). This suggests that the observed phage-induced fitness cost is not due to autoimmunity. Similarly, deep sequencing of bacterial populations that evolved phage resistance against DMS3vir showed that self-targeting with either zero (n = 17) or a single mismatch (n = 18) to the host-genome spacers were very rare (0.16% of all unique spacers). However, one spacer was found to target both the phage and host genome with 100% sequence identity (hypothetical protein YP_950448.1 and DUF1804 family protein respectively). While this spacer was present in two populations, it was only observed at the first timepoint sampled (day 2), consistent with the idea that there is strong selection against a self-targeting spacer. Our results therefore suggest that although self-targeting is likely to be deleterious, it is insufficient to explain the initial invasion of SM clones.

To explore whether expression of either host or phage genes following phage infection caused the phage-induced cost of CRISPR-based immunity, we performed RNAseq at 35, 60 and 120 minutes post-infection of a CRISPR immune clone that carries 2 spacers targeting DMS3vir (BIM2). BIM2 shows complete immunity to DMS3vir and no phage particles are produced following infection (data not shown; also see Landsberger *et al.* 2018). Differential expression analysis found no evidence that CRISPR-Cas expression is enhanced following infection (Fig. S5). Of all CRISPR-Cas genes, only Cas1 showed a significant difference, with slightly lower expression in infected BIM2 populations relative to controls (FDR adjusted p-value = 0.017, log2 fold change = - 0.375). Finally, to test whether we could detect phage gene expression in the infected BIM2 populations we mapped reads to the DMS3vir genome. This revealed significant levels of genome-wide phage gene expression in the BIM2 populations, despite their CRISPR-based immunity. In total, phage expression in the BIM2 populations was around 5-fold lower compared to infected WT populations (Fig. S6). Amongst the expressed phage genes, we identified very high levels of anti-CRISPR (*acr*) and associated repressor (*aca*) expression (Fig. S6), which has previously been reported to be amongst the most strongly expressed genes on the phage DMS3vir genome (Stanley *et al.* 2019), and to a lesser degree, many other genes (Fig S7). This *acr* protein is specific for type I-E CRISPR-Cas systems and does not impact the type I-F CRISPR-Cas system encoded by PA14. Collectively, these data show that the phage is capable of expressing its genes prior to CRISPR mediated cleavage.

To test whether expression of the *acr* or *aca* genes might be responsible for the phage-induced fitness cost in CRISPR-immune bacteria, we next competed the BIM2 strain and the surface mutant in the presence of either WT DMS3vir, an *acr* deletion mutant, and a mutant carrying a deletion of the entire *acr* operon. This revealed a similar fitness cost regardless of the phage genotype (GLM, effect of genotype on relative fitness of BIM vs SM, F_1,50_ = 2.15, p = 0.12). However, given that many other phage genes are also expressed, albeit at lower levels, the induced cost may well be due to expression of one or more other phage genes. We hypothesized that expression of the protease I gene might be particularly costly, since proteolytic activity could conceivably cause cytotoxicity in the cell. This essential gene is located immediately downstream of the *acr* and *aca* genes, and was found to be expressed both in this experiment as well as an independent RNAseq experiment using nanopore sequencing (data not shown). Given that a protease I deletion mutant would not be viable, we cloned and expressed the protease gene in WT *P. aeruginosa* to measure the cost of expression of this gene for the host. This showed that protease I expression reduced cell growth rates by approximately 13% relative to an empty vector control (GLM, F_1, 46_ = 28.72, p < 0.001). Given that this is just one of the many genes expressed by the phage during infection it seems likely that phage gene expression prior to clearance of the infection is a source of the observed phage-induced cost of CRISPR-based immunity. However, we note that there may be additional unknown factors that contribute to an inducible-cost.

## Discussion

Here we demonstrate that bacteria are rapidly immunised following phage infection and the population is driving the phage to extinction. However, the success of this immunisation is transient and successive re-introduction of the phage leads to stable coexistence between bacteria and phages. This is due to the invasion of bacteria with SM resistance as well as sensitive bacteria, which is caused by both an inducible fitness cost of CRISPR-based immunity and the loss of spacers. We attribute the inducible cost to the phage gene expression we observe which likely occurs prior to clearance of the infection by the CRISPR immune system. Immune loss has been shown to result in the stable coexistence of bacteria and phages (Weissman *et al* 2018.; Chaudhry *et al.* 2018) and the loss of spacers, resulting in a loss of immunity, is highly consistent the ecological dynamics we observe.

In a broader context, parasites are often invoked as a selective force for maintaining host diversity through negative frequency dependent selection (Koskella & Lively 2009). However, the effectiveness of CRISPR-Cas in driving phage extinct shortly after the onset of the phage epidemic (van Houte *et al.* 2016), removes this key mechanism for maintaining host spacer diversity. Indeed, we observed that spacer diversity decreased in all treatments (Fig. 3). This rapid decline in CRISPR diversity also helps to explain the observed conservation of the trailer-end of CRISPR loci (Tyson & Banfield 2007), although other ecological and evolutionary processes, such as selective sweeps of multi-phage resistant CRISPR clones (Weinberger *et al.* 2012), will also be important in natural communities. Interestingly, we find that surface modifications that confer phage resistance in this environment are not associated with a measurable trade-off in competitive fitness in the absence of phage. Although the variation in the genetic basis of the SM phenotype (SNPs and large deletions) suggests these mutations may vary in their fitness consequences in more complex environments. For example, increasing the environmental complexity, in the form of additional competitor species has been shown to amplify the associated fitness trade-off (Alseth *et al.* 2019). Therefore, understanding these dynamics across a range of environments will be of importance for future work.

The loss of spacer diversity over time is particularly relevant in the context of phage therapy, which is experiencing renewed clinical interest (Dedrick *et al.* 2019). Long-lasting resistance mechanisms may severely hamper therapeutic usage of phages, however this work shows that one can take advantage of the loss of population-level CRISPR immunity over time and design well-timed re-infection schemes. Clinical trials where burn wound patients are treated with cocktails of *Escherichia coli* and *P. aeruginosa* phages are ongoing (Rose *et al.* 2014) and experiments with murine models have demonstrated that phage therapy can provide highly effective treatment of *P. aeruginosa* infections (Waters *et al.* 2017; Wright *et al.* 2009; Kingwell 2015; Roach *et al.* 2017). This work also highlights an important potential issue associated with long-term information storage in CRISPR loci of bacterial populations (Shipman *et al.* 2017; Shipman *et al.* 2016), since unique spacer sequences will inevitably be lost over time.

## Materials and Methods

### Bacterial strains and phages

The previously described *P. aeruginosa* strains UCBPP-PA14 and the isogenic mutant *csy3::LacZ* and phage DMS3vir have been previously described (Cady *et al.* 2012) and were used throughout this study.

### Evolution experiments

Evolution experiments were performed in 6 replicates by inoculating 6 ml M9 supplemented with 0.2% glucose or LB with approximately 10^6^ bacteria from fresh overnight cultures of the WT strain and adding 10^7^ pfu of DMS3vir, followed by incubation at 37 °C while shaking at 180 rpm. Cultures were transferred daily 1:100 for 30 days. Reinfections with ancestral phage (10^7^ pfu) were performed as follows: a) bacterial cultures in the single infection treatment were only infected at t=0 b) bacterial cultures in the 5 day re-infection treatment were infected at t=0, 5, 10, 15, 20 and 25 dpi c) bacterial cultures in the 10 day re-infection treatment were infected at t=0, 10 and 20 dpi d) bacterial cultures in the daily infection treatment were infected at each daily transfer.

### Phage extraction and titration

Every second transfer, samples were taken just before transferring of the cultures into fresh medium. Phage was isolated from these samples using chloroform extractions. Next, phage was titrated by spotting serial dilutions of phage in M9 salts on a lawn of *P. aeruginosa* strain UCBPP-PA14 *csy3::LacZ* bacteria for quantification.

### Deep sequencing analysis

Full bacterial genomic DNA was isolated using the Qiagen QIAmp DNA mini kit as per the manufacturer’s protocols. A PCR amplification was performed for both CRISPR arrays (CRISPR1 and CRISPR2, see table S2). PCR reactions contained 5 μl DreamTaq master mix (ThermoScientific, UK), 0.5 μl forward primer, 0.5 μl reverse primer, 1.5 μl MiliQ water, 0.5 μl DMSO, 2 μl template DNA. Sample purity was determined by NanoDrop and DNA concentrations were quantified done using a Qubit fluorometer (ThermoFisher, UK).

### CRISPR array sequencing protocol

Two separate CRISPR primer (CRISPR1 and CRISPR2 locus) pairs were designed for two first round PCRs. 2 µl of DNA entered a first-round of PCR. The primer design incorporates a recognition sequence to allow a secondary nested PCR process. Samples were first purified with Ampure SPRI Beads before entering the second PCR performed to incorporate Illumina adapter sequences. Samples were purified using Ampure SPRI Beads before being quantified using Qubit and assessed using the Fragment Analyzer. Successfully generated amplicon libraries were taken forward and pooled in equimolar amounts, then size selected on a Pippin prep using a range of 180-600bps. The quantity and quality of each pool was assessed by Bioanalyzer and subsequently by qPCR using the Illumina Library Quantification Kit from Kapa on a Roche Light Cycler LC480II according to manufacturer’s instructions. Template DNA was denatured according to the protocol described in the Illumina cBot User guide and loaded at 12.5 pM concentration. To help balance the complexity of the amplicon library 15% PhiX was spiked in. The sequencing of each pool was carried out on one lane of an Illumina MiSeq, at 2×250 bp paired-end sequencing with v2 chemistry.

### Bioinformatics analysis

#### Sequence Quality Control

Base-calling and de-multiplexing of indexed reads was performed by CASAVA version 1.8.2 (Illumina) to produce 97 samples from each of the 2 lanes of sequence data. FASTQ files were trimmed to remove Illumina adapter sequences using Cutadapt version 1.2.1 (Martin 2011). The option “-O 3” was set, so the 3’ end of any reads which matched the adapter sequence over at least 3 bp was trimmed off. The reads were further trimmed to remove low quality bases, using Sickle version 1.200 with a minimum window quality score of 20. After trimming, reads shorter than 10 bp were removed. The raw reads were subjected to a Cutadapt trimming step to remove PCR primer sequences that could potentially introduce an artificial level of complexity in the samples. To improve base quality in both read pairs, sequencing errors were corrected in both forward and reverse reads using the error-correct module within SPAdes sequence assembler, version 3.1.0 (Bankevich *et al.* 2012). Read pairs were aligned to produce a single sequence for each pair that would entirely span the amplicon using PEAR (version 0.9.10; (Zhang *et al.* 2014)). Additionally, sequences with uncalled bases (Ns) were removed. To remove sequences originating from potential PCR primer dimers or from any spurious amplification events, a size selection was applied to each merged sequence set, respectively between 30-140bp for CRISPR1 and 70-500bp for CRISPR2. Fragmented PhiX phage genome was added to the sequence library in order to increase the sequence complexity. To remove any ‘bleed through’ of PhiX sequences, each sample was compared with the complete PhiX sequence (GenBank gi9626372) using BLASTN (Altschul *et al.* 1990). Sequences matching PhiX (E-value < 10-5) were filtered out of the dataset.

#### Clustering and diversity metrics

For each dataset, any sequences passing the filters (from any sample) were merged into a single file. This final sequence file, plus its own metadata file describing each sample, was used for the analysis by using a custom pipeline based on QIIME 1.9.0 (Caporaso *et al.* 2010). Clusters were defined using SWARM (Mahé *et al.* 2014), using the strictest (default) parameters. This tool aggregates a sequence to a cluster if the sequence shows similarity with any of the sequences already present in that cluster. Importantly, the similarity threshold is not fixed but defined within the dataset. A minimum cluster size filter is applied to retain clusters containing at least 2 sequences and potential chimeric-sequences due to PCR events were discarded as well. To calculate the abundance of each cluster, sequences were then aligned on the centroid sequence identified for each clusters, using a minimum similarity threshold of 99% for the entire length of the sequence using the ‘usearch_global’ function in VSEARCH.

The sequencing depth of all samples was explored using the ‘Chao 1’ (Chao, 1984) richness index plotted as a rarefaction curve. Counts in the cluster abundance tables were repeatedly sub-sampled (rarefied; 33 repetitions) at sampling depths of 1000, 12000, 22000, … 150000. The average Chao1 value obtained by repeating the test 33 times is assigned as alpha-diversity at that specific number of reads for that sample implemented in Qiime. Because all samples reached a clear asymptote, i.e. no samples were under-sampled with regards to spacer diversity, rarefaction was not applied. An abundance table for each locus was used to estimate the richness and evenness of the samples using the following estimators: total observed sequence variants, Shannon, Simpson, Simpson evenness again conducted with Qiime.

#### Whole-genome sequencing

Populations from T12 were plated and 12 resulting colonies were screened for their phage resistance phenotype, as described above. An additional control of the ancestral PA14 laboratory stock was included. Each colony was then suspended in 100 ul H_2_0 and streaked across an LB agar plate. Cells were scraped from these plates and added to bead beating tubes prior to shipping to the MicrobesNG service (Birmingham UK). Three beads were washed with extraction buffer containing lysozyme and RNase A, incubated for 25 min at 37 C. Proteinase K and RNaseA were added and incubated for 5 min at 65 C. Genomic DNA was purified using an equal volume of SPRI beads and resuspended in EB buffer. DNA was quantified in triplicates with the Quantit dsDNA HS assay in an Ependorff AF2200 plate reader. Genomic DNA libraries were prepared using Nextera XT Library Prep Kit (Illumina, San Diego, USA) following the manufacturer’s protocol with the following modifications: two nanograms of DNA instead of one were used as input, and PCR elongation time was increased to 1 min from 30 seconds. DNA quantification and library preparation were carried out on a Hamilton Microlab STAR automated liquid handling system. Pooled libraries were quantified using the Kapa Biosystems Library Quantification Kit for Illumina on a Roche light cycler 96 qPCR machine. Libraries were sequenced on the Illumina HiSeq using a 250bp paired end protocol. Reads were adapter trimmed using Trimmomatic 0.30 with a sliding window quality cutoff of Q15 (Bolger *et al.* 2014). The ancestral control was initially mapped to the PA14 reference genome (NC_008463) using the Breseq pipeline (version 0.32.0, (Deatherage & Barrick 2014). The reference genome was then modified to match the identified mutations (to account for divergence between the laboratory strain and original reference genome). The remaining samples were then mapped against this modified reference and alignments were used for identifying mutations.

### Immunity and resistance profiling

Bacterial immunity against the ancestral phage was determined as described before (Westra *et al.* 2015; van Houte *et al.* 2016; Chabas *et al.* 2018) by streaking individual clones (24 clones per sample) through ancestral phage DMS3vir and phage DMS3vir carrying the anti-CRISPR F1 (*acrIF1*) gene (Bondy-Denomy *et al.* 2012). Bacterial clones sensitive to both phages were scored as “sensitive”, those resistant to DMS3vir but sensitive to DMS3vir+AcrF1 were scored as “CRISPR immune”, and bacterial clones resistant to both phages were scored as “surface mutants”. CRISPR-Cas-mediated immunity was further confirmed by PCR using primers CTAAGCCTTGTACGAAGTCTC and CGCCGAAGGCCAGCGCGCCGGTG (for CRISPR 1) and GCCGTCCAGAAGTCACCACCCG and TCAGCAAGTTACGAGACCTCG (for CRISPR 2). Surface modification was further confirmed on the basis of colony morphology (since phage DMS3vir is pilus-specific, surface mutants have motility defects, resulting in a modified colony morphology) and a lack of new CRISPR spacers. From these analyses fractions of each phenotype (sensitive, CRISPR immune, surface mutant) were calculated for each replicate experiment.

### Statistical analyses

In most cases general linear models were used using the appropriate error structure and model residuals were assessed for model fit. Significance was determined following stepwise deletion of terms or through Tukey post-hoc testing with adjustment for multiple comparisons. General linear mixed-effect models were used for the competition assays, with population specified as a random effect, due to the non-independence of clones isolated from the same populations. Mixed effect modelling was carried out using the ‘lme4’ package with post-hoc testing conducted with the ‘emmeans’ package. For gene expression analysis, data were analysed with the DESeq2 pipeline using a parametric fit for the dispersion (Love *et al.* 2014). Statistical analyses were carried out in Rstudio using R version 3.5.2.

### Gene expression profiling

Ten replicate cultures of PA14 BIM2 and 5 of WT were grown in microcosms of 6 mL LB media and standardized to an optical density of 0.5 OD600 (approx. 2 × 10^9 CFU / mL). Five cultures of BIM2 and 5 cultures of WT were inoculated with 8 × 10^9 PFU (in 500 uL) DMS3vir (MOI 0.5) and 5 cultures of BIM2 remained uninoculated as controls. After 35 minutes, 1 hour and 2 hours, 1.5 mL of cells were pelleted and snap-frozen at – 80 C. For RNA extraction, 1 mL of TRIzol reagent was added immediately after removal from freezer. BCP was used as a phase separator followed by PureLink RNA extraction (Invitrogen) that included a DNAse treatment step. Sample quantities, purities and size distribution were quantifed by Qubit, nanodrop and TapeStation respectively. 150bp libraries were prepared with the TruSeq directional kit and sequenced on an Illumina NovaSeq at Exeter University sequencing centre.

The resulting reads were quality filtered using Sickle (default settings, version 1.33) with approx. 6 million read pairs per sample remaining. These were subsequently mapped to the PA14 and DMS3vir reference genomes with bwa (default settings, version 0.7.17). The resulting read alignments were assigned to genomic features and counted using HTSeq (version 0.11.2, (Anders *et al.* 2015) using ‘union’ mode, non-unique setting set to ‘none’ and the reverse stranded option. Differential expression analysis was conducted in R using the DESeq2 package (Love *et al.* 2014).

### ProteaseI expression

The proteaseI gene from DMS3vir was cloned into the pHERD30T vector under control of an arabinose inducible promoter. Primers used for cloning are available in table S3. Colonies were selected using a gentamycin marker and blue/white colony screening in a background of dH5-alpha cells (New England Biolabs, UK). The resulting construct was transformed into PA14 and the clone used for assays was verified by Sanger sequencing (University of Sheffield, UK). Optical density was recorded during a 24-hour growth curve at 37 C in LB supplemented with 1% arabinose. Growth curves of the clone expressing proteaseI and an empty vector control were recorded. Growth rate during log-phase was extracted from the growth curves in R (version 3.5.2).

### Competition assays

For each population, a single clone of each phenotype (CRISPR, SM, Sens.) was picked (where possible) and competed against a LacZ marked reference strain. Populations were serially transferred with a 1:100 dilution in M9 media supplemented with 0.2% glucose. Populations were plated at T = 0 and after 3 days on LB agar supplemeted with 30 μg of X-gal to determine the relative frequencies of the clone and the reference. Selection coefficients were determined as described in (van Houte *et al.* 2016).

### Data availability

Sequence data is available from the European Nucleotide Archive under study number PRJEB31514.

## Author contributions

E.R.W., S.v.H., S.M. and A.B. designed experiments. L.C. and S.M. performed the experiments under supervision of S.v.H. and E.R.W. E.O. prepared samples for sequencing. S.M., L.L., A.Biswas. performed sequencing data analyses. E.R.W. and S.M. wrote the manuscript.

## Acknowledgements

E.R.W. and A.B. acknowledge the Natural Environment Research Council, and the Biotechnology and Biological Sciences Research Council for funding. E.R.W further acknowledges the Wellcome Trust and the European Research Council for funding. A.B further acknowledges the Royal Society, the Leverhulme Trust, and the AXA research fund for funding. SVH acknowledges funding from BBSRC (grant number BB/R010781/1). Whole genome sequencing was provided by MicrobesNG (http://www.microbesng.uk) which is supported by the BBSRC (grant number BB/L024209/1). RNA-sequencing was conducted at the University of Exeter sequencing centre which is supported by the Wellcome Trust.

## Supplementary Information

**Figure S1.**
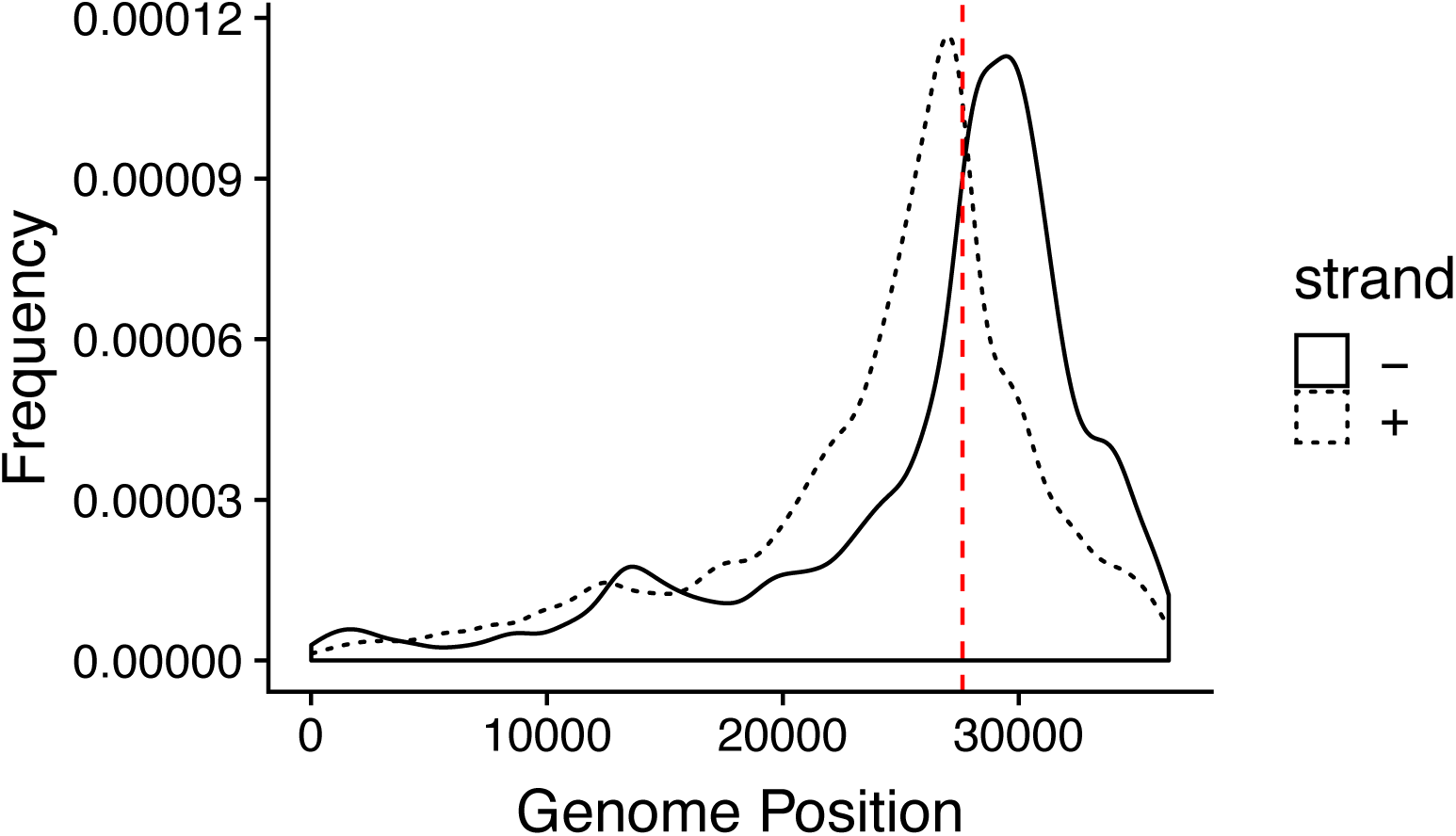
Locations of extracted spacers from the CRISPR arrays when mapped to the DMS3vir genome. Red line denotes the location of the priming spacer (CRISPR2 spacer 1).

**Figure S2.**
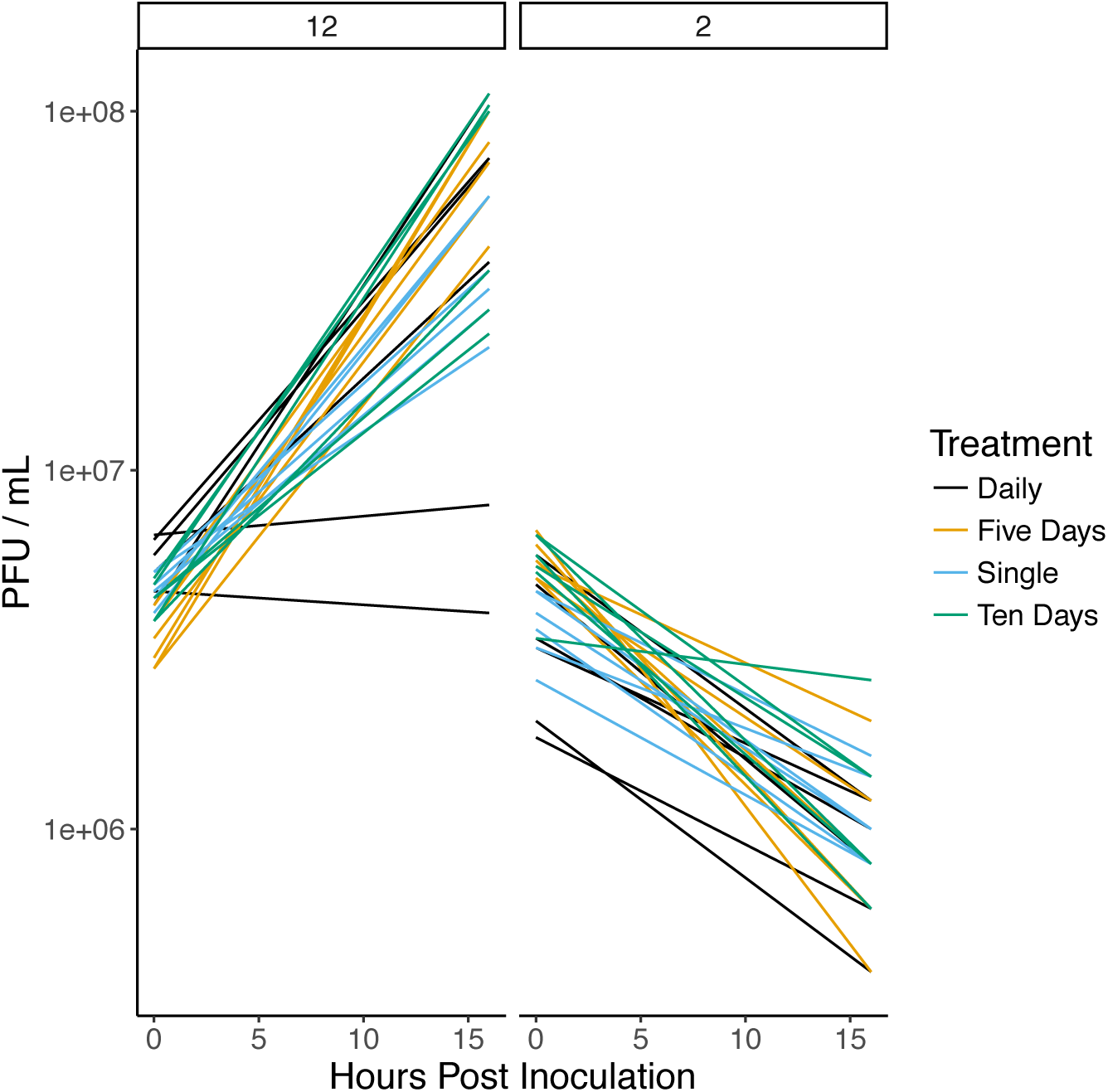
Changes in phage titres upon infection with 10^7^ PFU DMS3vir of populations isolated from day 12 (left panel, n = 24) or day 2 (right panel, n = 24) from each of the different reinfection regimes. Colours denote the reinfection regimes where the bacterial populations were derived from: bacterial populations isolated from the single infection regime are shown in blue, those from re-infection every five days are shown in yellow, reinfection every 10 days in green, and daily infection in black.

**Fig. S3.**
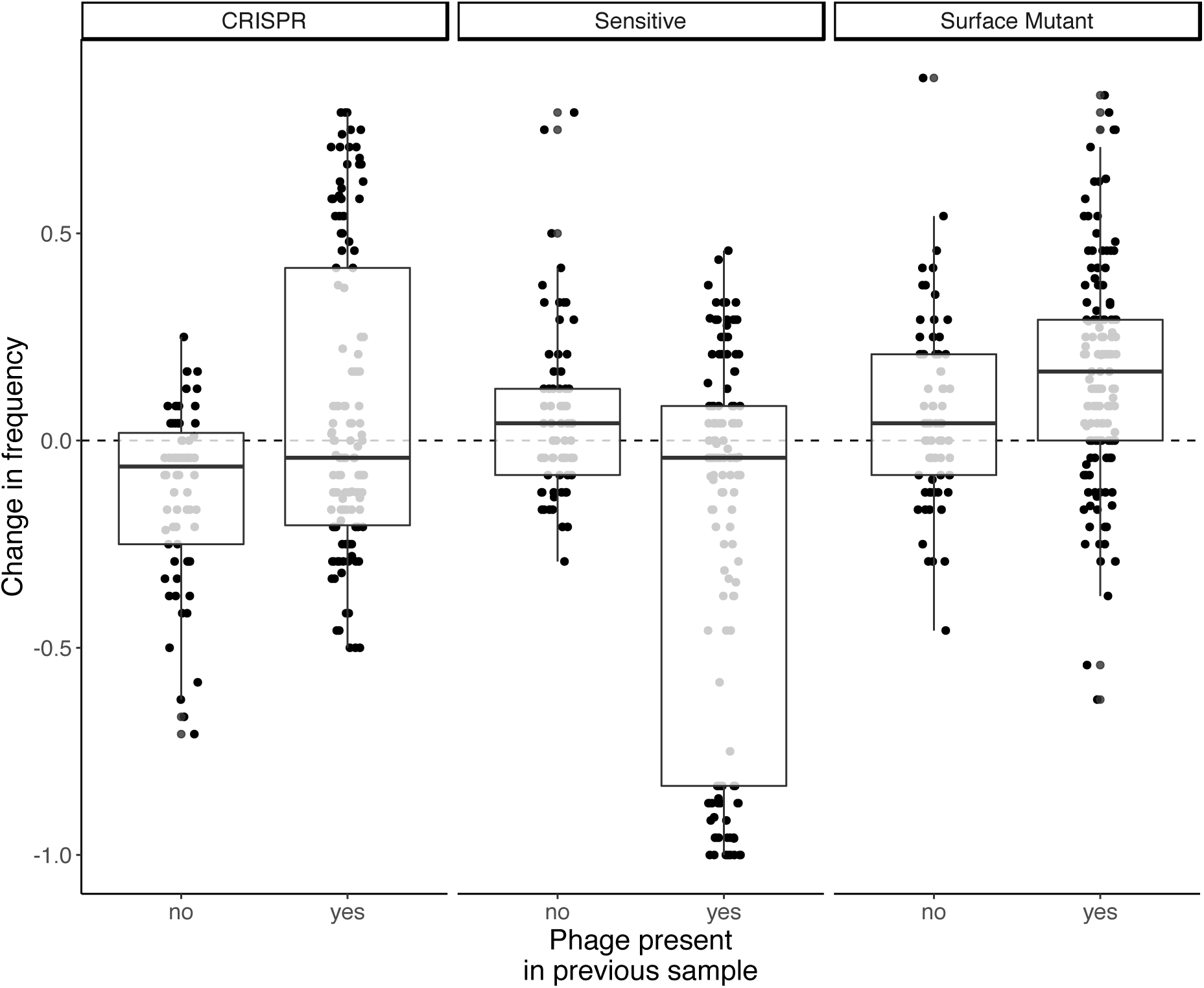
Change in frequency of each phenotype between sampling points during evolution experiment split by presence (n = 402) or absence (n = 215) of phage in the previous sample. Limit of detection for phage presence is 200 PFU per ml. Boxplots show the median, interquartile range and minimum and maximum values.

**Figure S4.**
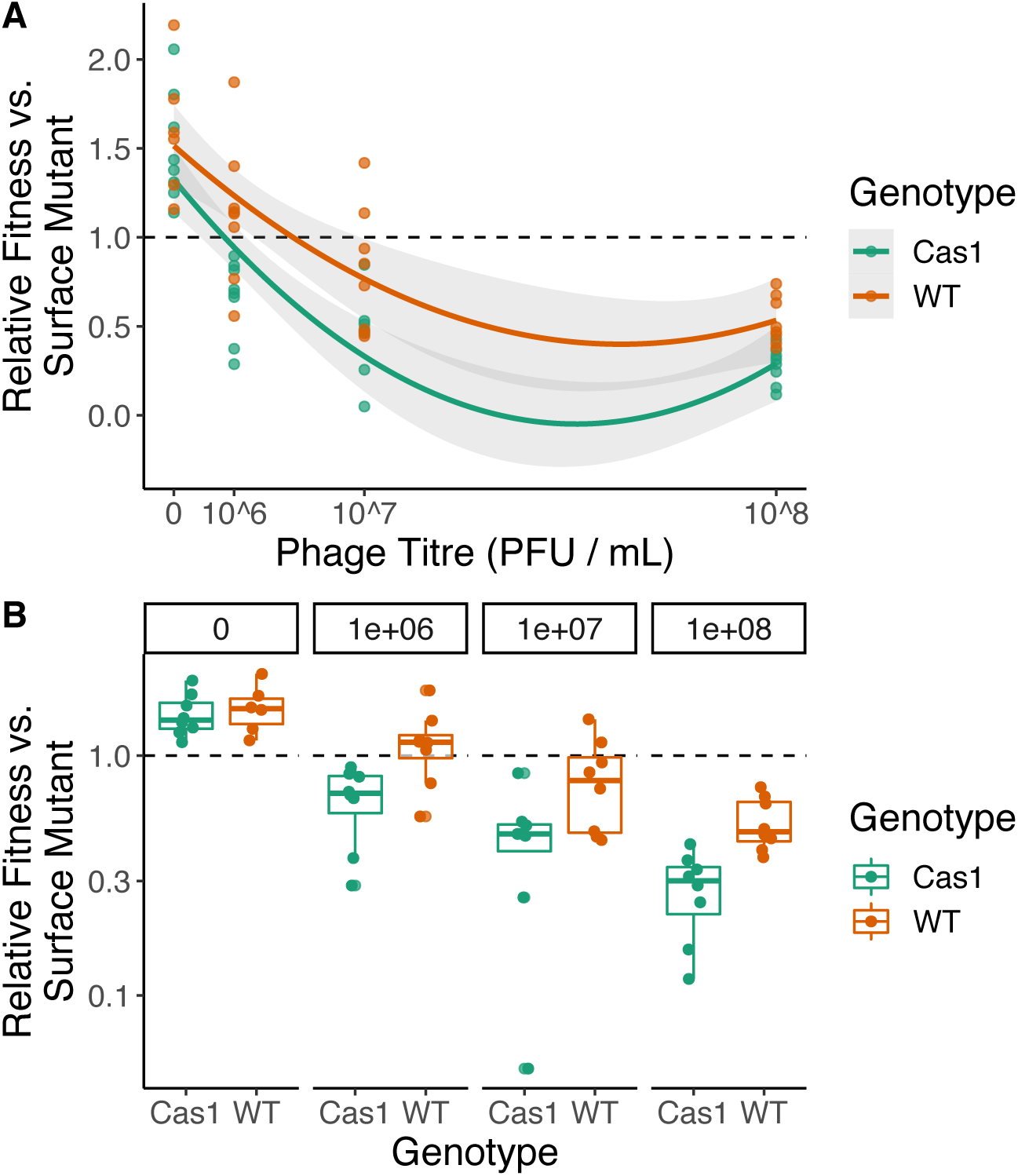
Relative fitness of bacterial populations with CRISPR-mediated immunity at 3 d.p.i. when competing with a surface mutant. Competitions were carried out between a marked surface resistant strain and either the WT PA14 (purple), a mutant PA14 lacking a functional *cas1* (orange) or a WT-derived strain carrying 2 additional spacers targeting phage DMS3mvir (BIM2; green). Experiments were carried out across a gradient of phage titres, as indicated. The phage used (DMS3mvir) is a mutant of DMS3vir that is targeted by WT PA14. N = 8 replicates per genotype per phage titre. A) Data presented with phage titre as a continuous variable. B) Data grouped by phage titre to better visualise differences within titres.

**Fig. S5.**
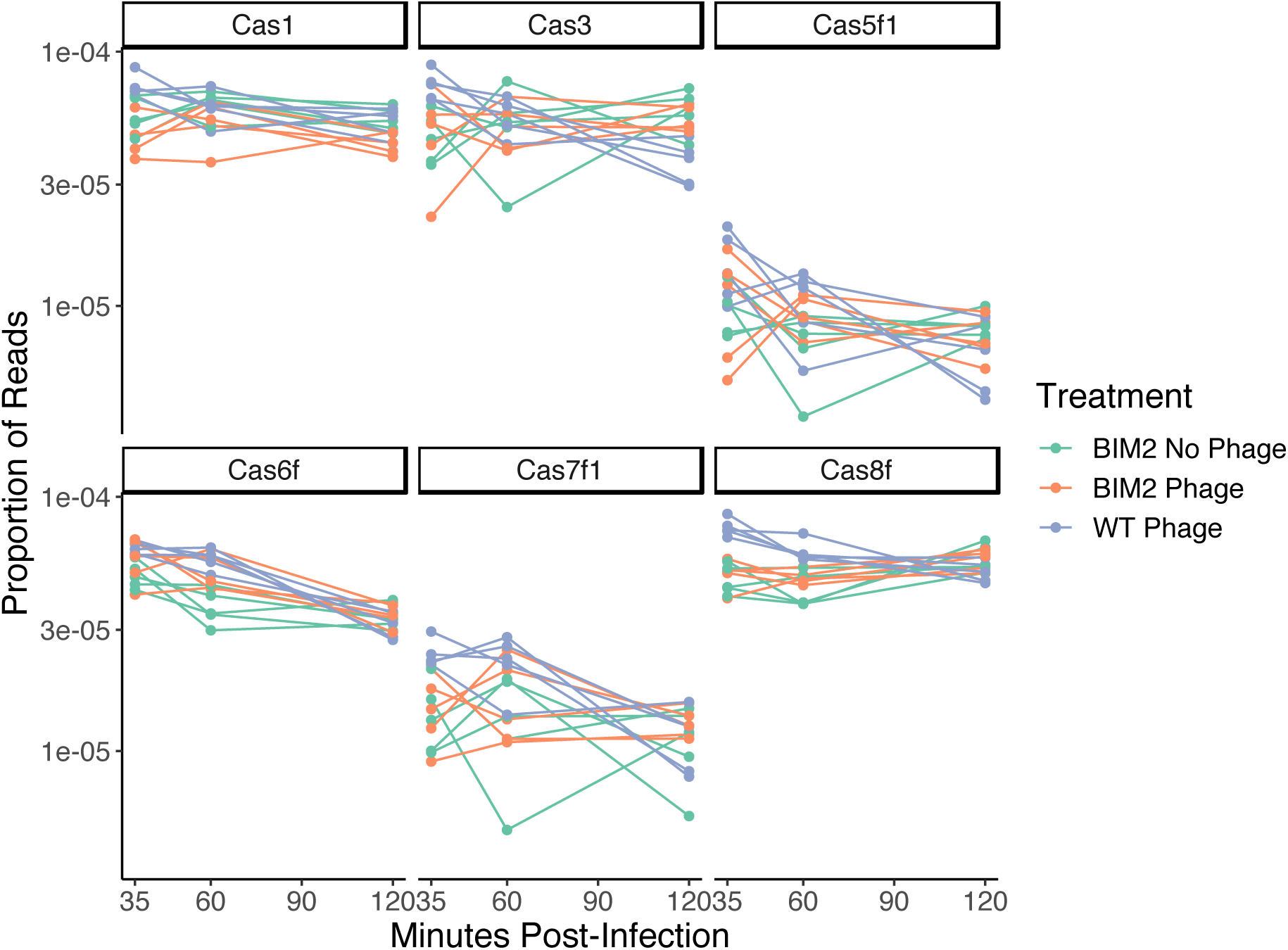
Expression of CRISPR associated genes following infection with 8 × 10^9 PFU DMS3vir (MOI 0.5) across PA14 WT (purple, n = 5), PA14 BIM2 (orange, n = 5) and PA14 BIM2 uninfected controls (green, n = 5). Samples taken 35, 60 and 120 minutes post-infection.

**Fig. S6.**
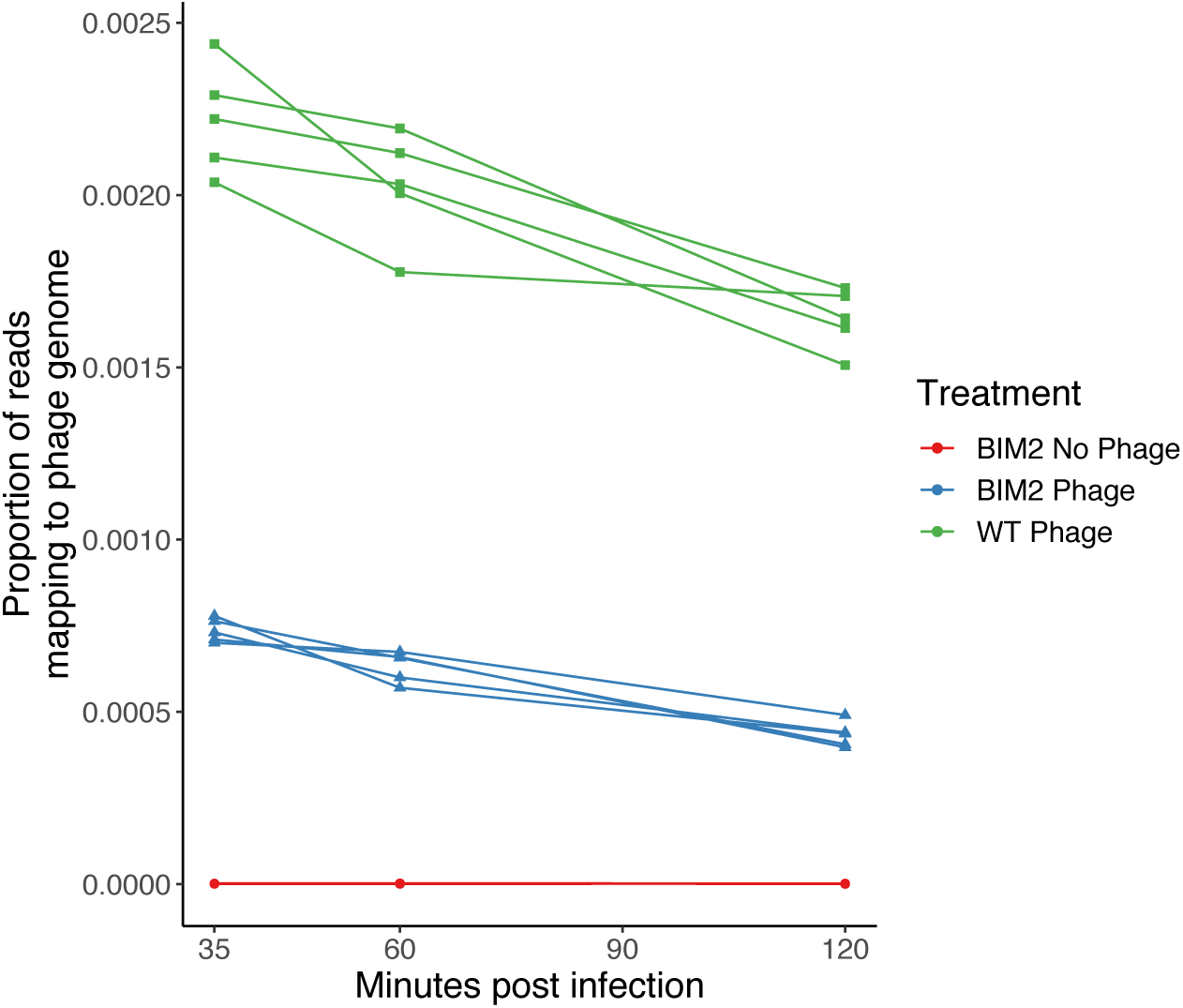
Total phage gene expression following infection with 8 × 10^9 PFU DMS3vir (MOI 0.5) across PA14 WT (green, n = 5), PA14 BIM2 (blue, n = 5) and PA14 BIM2 uninfected controls (red, n = 5). Samples taken 35, 60 and 120 minutes post-infection.

**Fig. S7.**
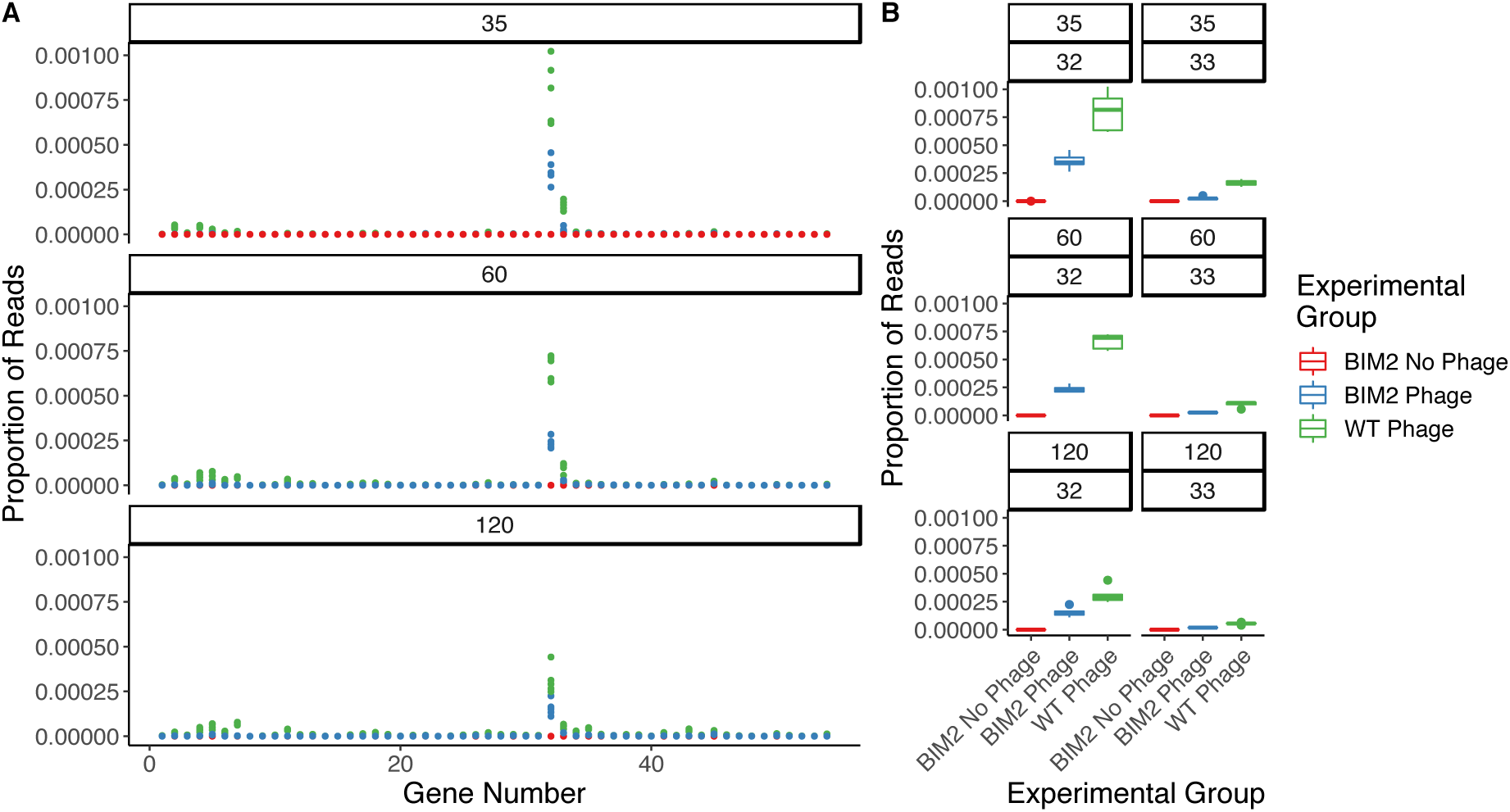
A) Phage gene expression of each annotated phage gene following infection with 8 × 10^9 PFU DMS3vir (MOI 0.5) across PA14 WT (green, n = 5), PA14 BIM2 (blue, n = 5) and PA14 BIM2 uninfected controls (red, n = 5) at 35, 60 and 120 minutes post-infection. B) Gene expression of the *acr* protein (32) and its *aca* repressor (33) across sampling times and treatments.

**Table S1.** Post-hoc comparisons of contrasts that compare relative fitness. Numbers in the contrast denote sampling time (day 4 or day 12) and the given phenotypes (CRISPR, SM or sensitive).

| Contrast | Estimate | SE | df | t.ratio | p-value |
| --- | --- | --- | --- | --- | --- |
| 4_CRISPR-12_CRISPR | -0.2773 | 0.0474 | 122.8124 | -5.8475 | **1e-04** |
| 4_CRISPR-4_Sens | -0.1115 | 0.0586 | 109.4472 | -1.9017 | 0.4067 |
| 4_CRISPR-12_Sens | -0.3887 | 0.0742 | 111.9035 | -5.2423 | **1e-04** |
| 4_CRISPR-4_SM | -0.1243 | 0.0516 | 96.3502 | -2.4093 | 0.1634 |
| 4_CRISPR-12_SM | -0.4015 | 0.0693 | 111.6448 | -5.7951 | **1e-04** |
| 12_CRISPR-4_Sens | 0.1658 | 0.0766 | 120.3732 | 2.1639 | 0.2625 |
| 12_CRISPR-12_Sens | -0.1115 | 0.0586 | 109.4472 | -1.9017 | 0.4067 |
| 12_CRISPR-4_SM | 0.153 | 0.0708 | 113.2484 | 2.1593 | 0.2652 |
| 12_CRISPR-12_SM | -0.1243 | 0.0516 | 96.3502 | -2.4093 | 0.1634 |
| 4_Sens-12_Sens | -0.2773 | 0.0474 | 122.8124 | -5.8475 | **1e-04** |
| 4_Sens-4_SM | -0.0128 | 0.058 | 108.8156 | -0.2211 | 0.9999 |
| 4_Sens-12_SM | -0.2901 | 0.0754 | 119.9723 | -3.8455 | **0.0026** |
| 12_Sens-4_SM | 0.2644 | 0.0744 | 112.3086 | 3.5541 | **0.0072** |
| 12_Sens-12_SM | -0.0128 | 0.058 | 108.8156 | -0.2211 | 0.9999 |
| 4_SM-12_SM | -0.2773 | 0.0474 | 122.8124 | -5.8475 | **1e-04** |

**Table S2.**
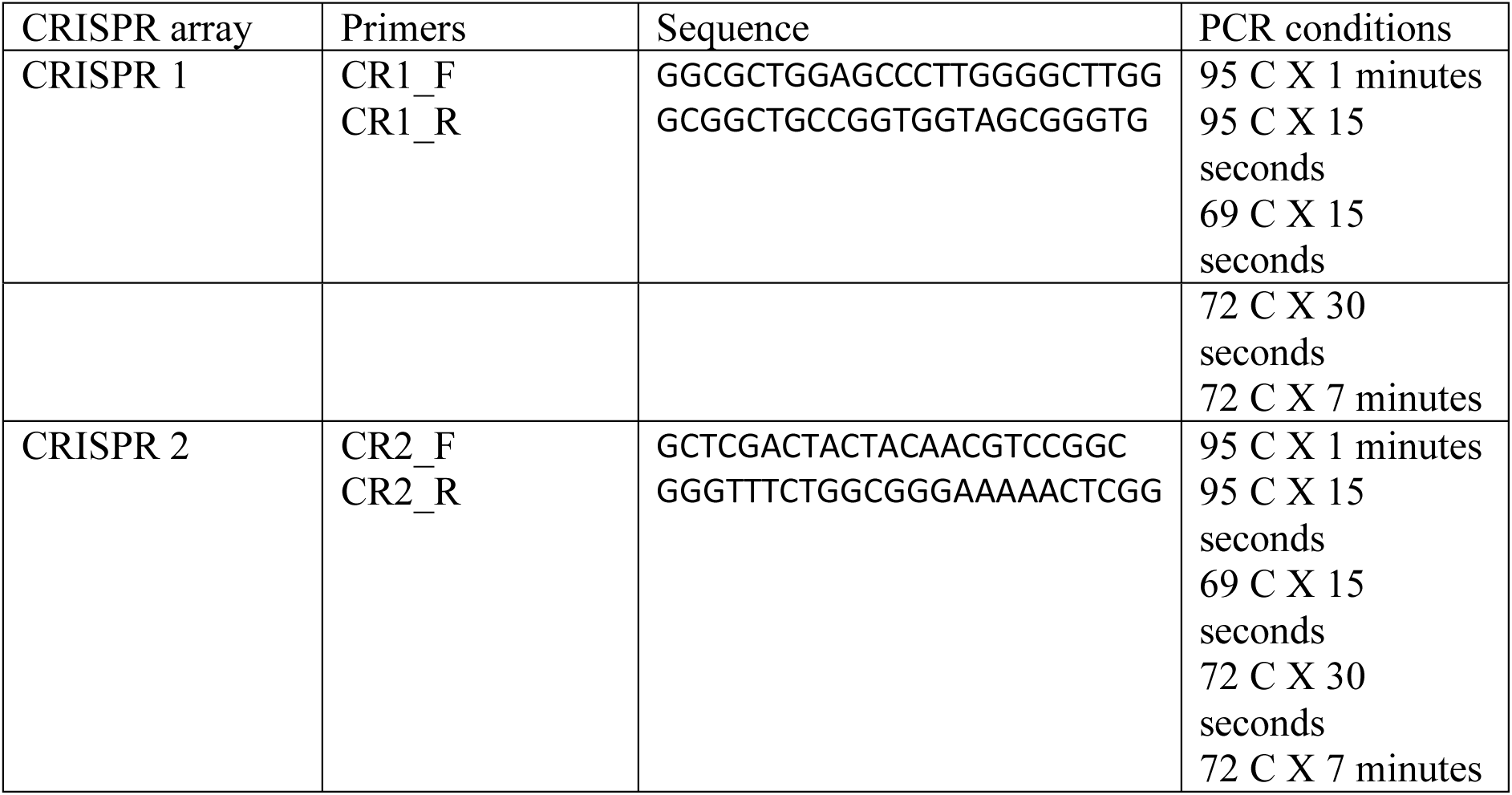
Primers and PCR conditions used for amplicon sequence analysis:

**Table S3.**
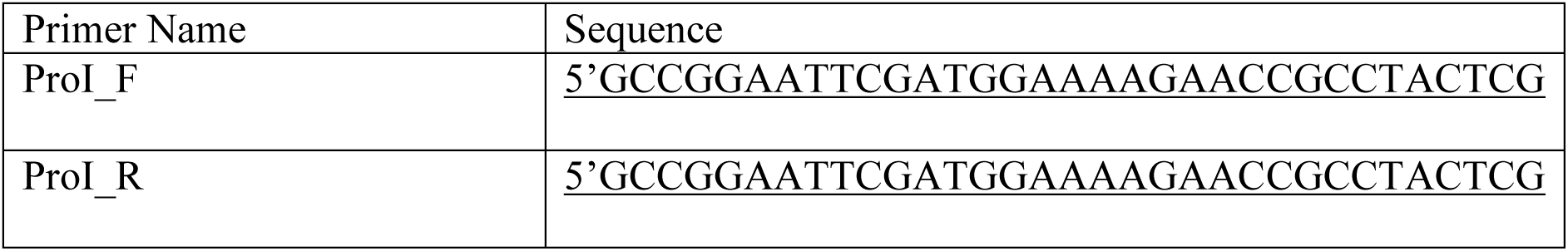
Primers for cloning proteaseI gene of DMS3vir

